# FGF21-FGFR4 signaling in cardiac myocytes promotes concentric cardiac hypertrophy in mouse models of diabetes

**DOI:** 10.1101/2021.09.16.460632

**Authors:** Christopher Yanucil, Dominik Kentrup, Xueyi Li, Alexander Grabner, Karla Schramm, Eliana C. Martinez, Jinliang Li, Isaac Campos, Brian Czaya, Kylie Heitman, David Westbrook, Adam Wende, Alexis Sloan, Johanna M. Roche, Alessia Fornoni, Michael S. Kapiloff, Christian Faul

## Abstract

Fibroblast growth factor (FGF) 21, a hormone that increases insulin sensitivity, has shown promise as a therapeutic agent to improve metabolic dysregulation. Here we report that FGF21 directly targets cardiac myocytes by binding β-klotho and FGF receptor (FGFR) 4. In combination with high glucose, FGF21 induces cardiac myocyte growth in width mediated by extracellular signal-regulated kinase 1/2 (ERK1/2) signaling. While short-term FGF21 elevation can be cardio-protective, we find that in type 2 diabetes (T2D) in mice, where serum FGF21 levels are elevated, FGFR4 activation induces concentric cardiac hypertrophy. As T2D patients are at risk for heart failure with preserved ejection fraction (HFpEF), we propose that induction of concentric hypertrophy by elevated FGF21-FGFR4 signaling constitutes a novel mechanism promoting T2D-associated HFpEF and that FGFR4 blockade might serve as a cardio-protective therapy in T2D. In addition, potential adverse cardiac effects of FGF21 mimetics currently in clinical trials should be investigated.

## Introduction

The global prevalence of type 2 diabetes (T2D) is growing rapidly and is expected to reach 300 million by 2025. Diabetes is a risk factor for congestive heart failure, and cardiovascular complications are the leading cause of diabetes-related morbidity and mortality (Jia et al., 2018). Diabetic cardiomyopathy is termed as heart failure in the absence of other comorbid conditions and is associated with both diastolic and systolic cardiac dysfunction, as well as pathological cardiac myocyte hypertrophy (Jia et al., 2018). Cardiac remodeling includes defects in calcium handling and myocyte contractility, altered myocyte metabolism, and an increase in oxidative stress and mitochondrial dysfunction, as well as induction of interstitial myocardial fibrosis. Over 40% of T2D patients have increased left ventricular mass and wall thickness, conferring additional risk of heart failure that is independent of the other features of metabolic syndrome (hypertension, hyperlipidemia, and obesity) often present in these patients (Aneja et al., 2008; Devereux et al., 2000; Virani et al., 2021). Importantly, hyperglycemia and the associated metabolic abnormalities in T2D are considered major factors promoting heart failure with preserved ejection fraction (HFpEF) (Mishra and Kass, 2021). An area of intense current research, HFpEF represents about of half of all patients with heart failure and is a syndrome for which there is currently no effective medical therapy. The present lack of HFpEF therapies is in part a consequence of our poor understanding of the molecular mechanisms inducing pathological cardiac remodeling in diabetic cardiomyopathy. Here we consider the potential role of FGF21 in the development of pathological concentric cardiac hypertrophy.

The fibroblast growth factor (FGF) family consists of 22 members that by activation of FGF receptor (FGFR) tyrosine kinases regulate cellular proliferation, survival, migration and differentiation (Geng et al., 2020). FGF19, FGF21 and FGF23, are distinguished by their low heparin binding affinity and their function as hormones acting on distant target organs (Degirolamo et al., 2016). As the endocrine FGFs bind heparin poorly, cellular activation requires klotho, a family of single-pass transmembrane proteins (α- and β-klotho) that serve as co-receptors to facilitate FGF-FGFR binding. The klotho co-receptors have a more restricted expression pattern than FGFRs, thereby conferring tissue-specificity to the actions of endocrine FGFs (Degirolamo et al., 2016).

Produced mainly by hepatocytes, FGF21 functions as a major regulator of metabolism and energy homeostasis during fasting conditions (Fisher and Maratos-Flier, 2016; Geng et al., 2020). For example, by binding FGFR1/β-klotho receptors in adipocytes, FGF21 induces glucose uptake and fatty acid storage (Owen et al., 2015). Accordingly, pharmacological administration of FGF21 can have beneficial effects in animal models, including a reduction in blood glucose, triglyceride and cholesterol levels, weight loss, and increased life span (Owen et al., 2015). In patients with T2D and obesity, however, markedly elevated serum FGF21 levels have been observed (An et al., 2012; Chavez et al., 2009; Mraz et al., 2009; Zhang et al., 2008), prompting the hypothesis that obesity is a FGF21-resistant metabolic state (Planavila et al., 2015b). Elevated serum FGF21 levels are also present in hypertension (Semba et al., 2013), dilated cardiomyopathy (Gu et al., 2020), coronary artery disease (Haberka et al., 2019; Lin et al., 2010; Shen et al., 2013), acute myocardial infarction (Sunaga et al., 2019), and atrial fibrillation (Han et al., 2015). Notably, FGF21 expression has been found to be increased in HFpEF patients with diastolic dysfunction (Chou et al., 2016). FGF21 appears to have direct effects on the heart, including cardio-protective actions of short-term FGF21 elevations during ischemia/reperfusion injury, β-adrenergic activation, or hypertension (Hu et al., 2018; Joki et al., 2015; Li et al., 2020b; Li et al., 2019; Liu et al., 2013; Patel et al., 2014; Planavila et al., 2013; Planavila et al., 2015a; Ruan et al., 2018; Sun et al., 2019).

Concentric cardiac hypertrophy and diastolic dysfunction are prominent features of diabetic cardiomyopathy and HFpEF (Mishra and Kass, 2021). Concentric cardiac hypertrophy is characterized by an increase in relative wall thickness (left ventricular wall thickness to internal diameter ratio) that is primarily a reflection of increased cardiac myocyte width, while, in contrast, eccentric cardiac hypertrophy is characterized by decreased relative wall thickness and relative myocyte lengthening. Concentric hypertrophy can be induced by activation of an extracellular signal-regulated kinase 1/2 (ERK1/2) - p90 ribosomal S6 kinase type 3 (RSK3) - serum response factor (SRF)-dependent gene expression regulatory pathway (Kehat et al., 2011; Li et al., 2020a; Passariello et al., 2016). It has been observed that in cardiac myocytes FGF21 activates ERK1/2 and, like ERK1/2 gene deletion, unstressed FGF21 knock-out mice exhibit eccentric cardiac hypertrophy (Planavila et al., 2013). Here we consider the novel hypothesis that, despite its ability to promote myocyte survival, the elevation of FGF21 in T2D and HFpEF is not beneficial, but deleterious, such that in the context of T2D elevated FGF21 promotes pathological concentric cardiac hypertrophy. Studies performed both *in vitro* and *in vivo* suggest that FGF21 is a key humoral mediator of the pathological remodeling in diabetic cardiomyopathy, FGF21-FGFR4 signaling constituting a novel therapeutic target for heart failure associated with T2D.

## Results

### FGF21 induces concentric cardiac hypertrophy in mice

To test for possible effects on cardiac structure and function by elevated FGF21 levels, a FGF21 transgenic mouse (FGF21-Tg) was utilized in which FGF21 is systemically elevated by expression in the liver under the control of the apolipoprotein E (ApoE) promoter (Inagaki et al., 2007). Consistent with the original report (Inagaki et al., 2007), at 24 weeks of age these mice had reduced body weight and blood glucose levels, and a 200-fold increase in serum FGF21 levels (Figure 1A-C). While the analysis of cardiac function in FGF21-Tg mice was confounded by their significantly reduced body weight, by echocardiography these mice exhibited a persistent elevation in ejection fraction and relative left ventricular (LV) hypertrophy, as indicated by increased LV mass to body weight ratio and relative wall thickness (“concentricity index”) (Khouri et al., 2010), consistent with the development of concentric cardiac hypertrophy (Figure 1D-F, Supplemental Figure 1A; Supplemental Table 1). Relative cardiac hypertrophy in FGF21-Tg mice was confirmed by gravimetric measurement of heart weight to body weight ratio (Figure 1G). Paradoxically, cross-sectional area of cardiac myocyte was not increased in FGF21-Tg mice, presumably lower in proportion to the decreased body weight (Figure 1H,I). Histological and gene expression analyses did not show evidence of cardiac fibrosis in FGF21-Tg mice (Supplemental Figure 2A-D). Taken together, overexpression of exogenous FGF21 in mice appeared to induce concentric cardiac hypertrophy.

**Figure 1.**
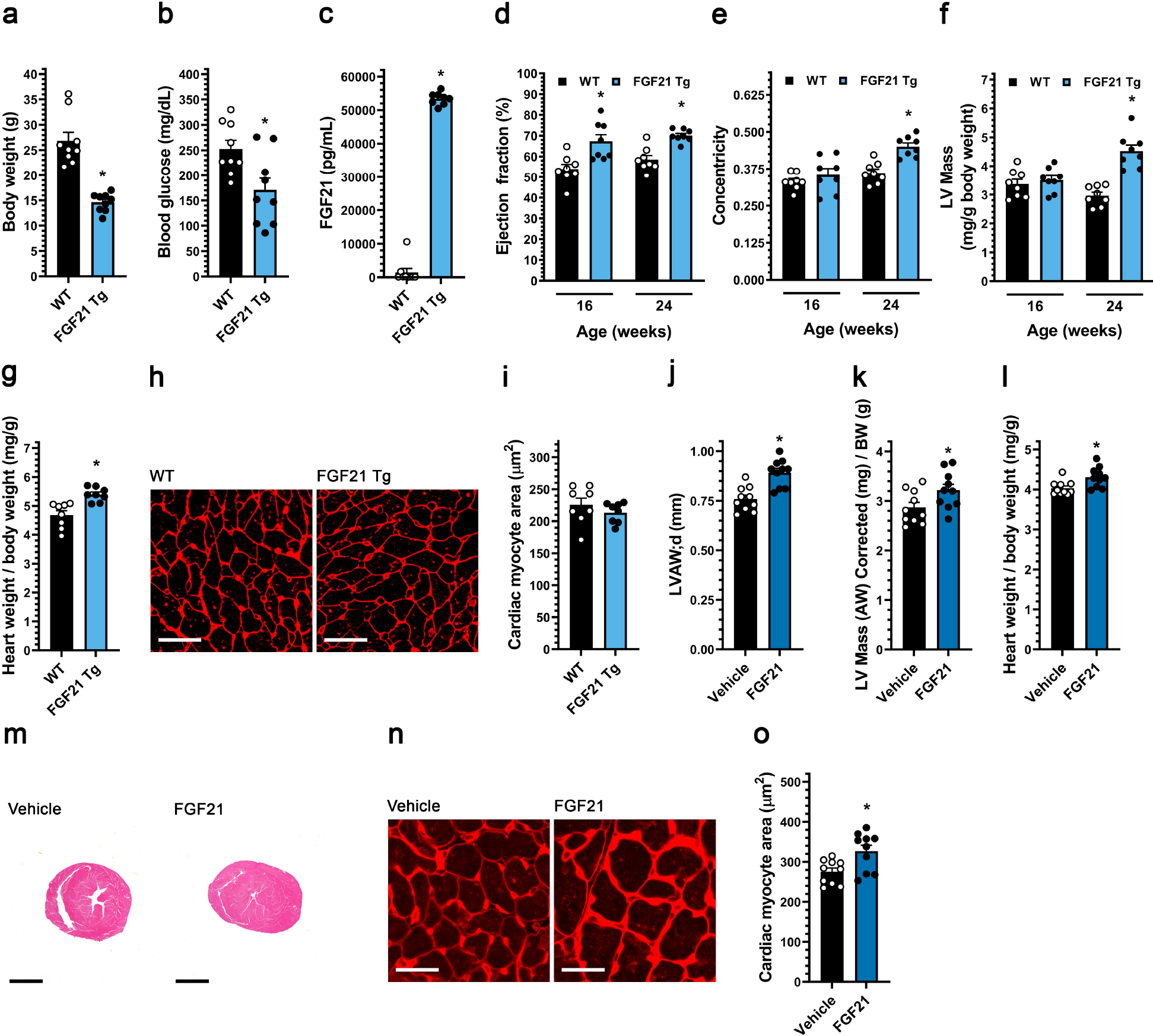
Systemic FGF21 elevation in mice induces concentric cardiac hypertrophy. (a-h) Analysis of FGF21 transgenic (Tg) mice and wild-type littermates at 24 weeks of age unless otherwise indicated. (a) Body weight, (b) blood glucose levels, and (c) serum FGF21 levels. (d) Ejection fraction, (e) concentricity, and (f) left ventricular (LV) mass, as determined by echocardiography at 16 and 24 weeks of age. (g) Heart weight/body weight ratio, (h-i) wheat germ agglutinin staining of LV tissue (scale bar = 25 µm) and cardiac myocyte cross-section area. (j-o) Wildtype mice were injected i.v. with FGF21 or vehicle for five consecutive days before analysis on the 6th day. (j) LV anterior wall thickness (diastole), and (k) LV mass/body weight ratio, as determined by echocardiography. (l) Gravimetric heart weight/body weight ratio. (m) hematoxylin and eosin-stained transverse heart sections (scale bar = 2 mm). (n-o) wheat germ agglutinin staining of LV tissue (scale bar = 25 µm) and cardiac myocyte cross-section area. Statistical significance was determined by 2-way ANOVA with post-hoc testing with Sidak’s multiple comparisons test (d-f), or by two-tailed t-test. All values are expressed as mean ± SEM. a, b) N=9; c) N=8-9; d-f) N=8, *p≤ 0.05 vs. WT of same age; g, i) N=8, *p≤ 0.05 vs. WT; j-l, o), N=10, *p≤ 0.05 vs. vehicle. For the complete set of echocardiography parameters see Supplemental Table 1 and 2.

To determine if FGF21 elevation could induce concentric cardiac hypertrophy in the absence of confounding effects on body weight, adult wildtype Balb/c mice were injected intravenously (i.v.) twice daily with 40 µg/kg of recombinant FGF21 protein or isotonic saline for five days, followed by echocardiography and tissue collection on the 6th day. Echocardiographic analysis revealed that FGF21 increased LV anterior wall thickness, LV mass and concentricity index (Figure 1J,K, Supplemental Figure 1B; Supplemental Table 2). Accordingly, gravimetric and histological analysis showed an elevation in indexed heart weight (Figure 1L,M) and in the cross-sectional area of individual cardiac myocytes (Figure 1N,O). These results demonstrate that elevated FGF21 can rapidly induce concentric cardiac hypertrophy in wildtype mice.

### FGF21 promotes the concentric hypertrophy of adult rat ventricular myocytes *in vitro*

Systemic FGF21 elevation *in vivo* resulted in concentric cardiac hypertrophy. To demonstrate that FGF21 can induce cardiac myocyte hypertrophy in a cell autonomous manner, primary adult rat ventricular myocytes (ARVM) were studied *in vitro*. Stimulation by 25 ng/ml FGF2, an FGF isoform known to induce myocyte hypertrophy (Corda et al., 1997), but not stimulation by FGF21 induced the hypertrophy of myocytes cultured in minimal medium (data not shown). Given that elevated FGF21 serum levels are associated with diabetic cardiomyopathy (Degirolamo et al., 2016), we considered that FGF21 might induce hypertrophy only in the presence of other stimuli, such as elevated glucose levels. Myocytes were cultured in the presence of 15.6 mM glucose, mimicking hyperglycemia of a serum level of 279 mg/dL and 3-fold the glucose concentration in standard minimal medium. While 48 hours of FGF21 stimulation did not induce myocyte hypertrophy, culture in medium containing elevated glucose significantly increased myocyte width 6%, while not affecting myocyte length, resulting in a significantly increased width:length ratio indicative of a mild “concentric” myocyte hypertrophy (Figure 2A-D). Remarkably, addition of FGF21 to medium containing high glucose further increased myocyte growth in width without affecting myocyte length, such that ARVM cultured in FGF21 and 15.6 mM glucose were 12% wider and had a width:length ratio 10% greater than control cells.

**Figure 2.**
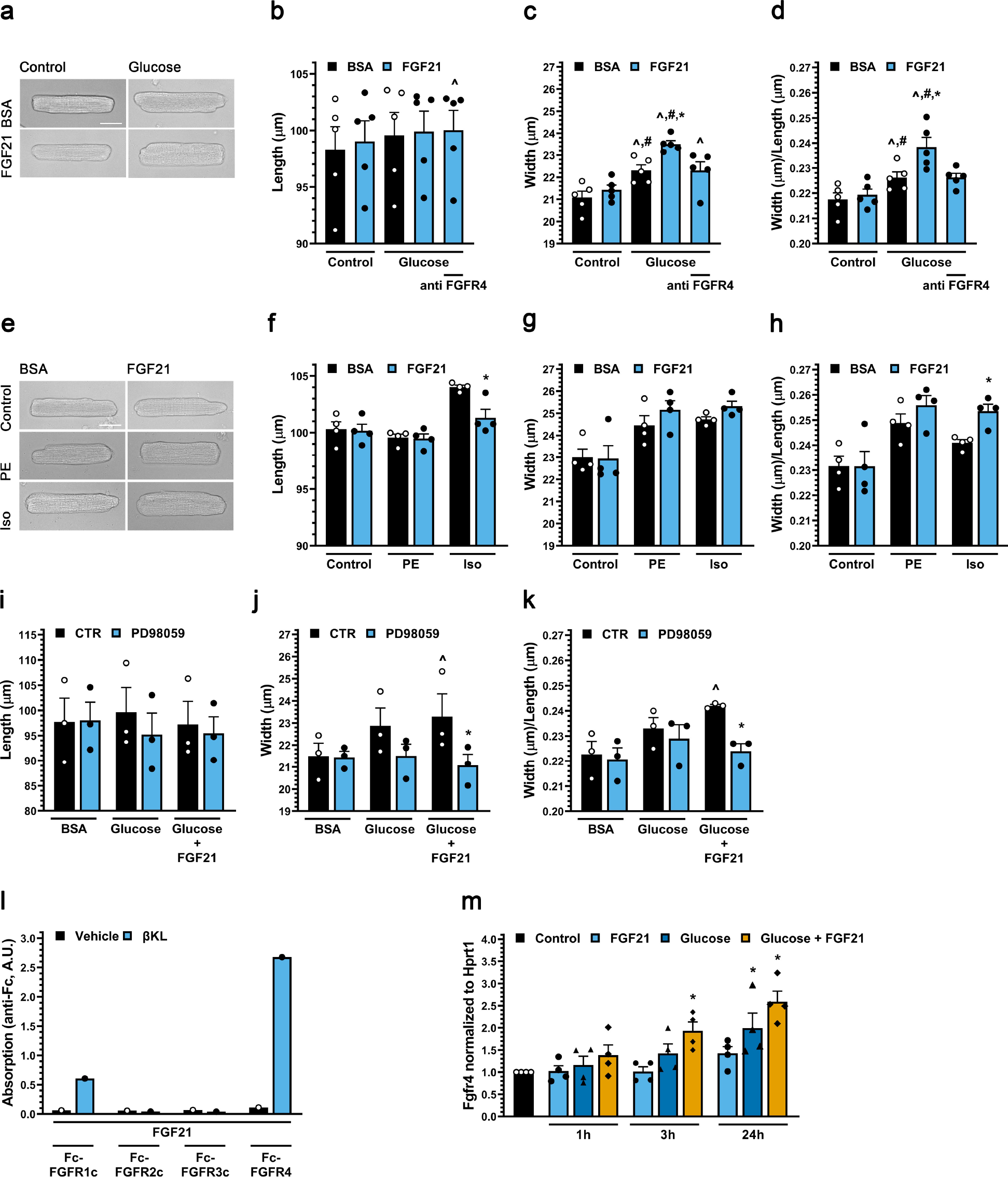
FGF21 induces hypertrophic growth of cultured cardiac myocytes in the presence of high glucose via FGFR4 and ERK1/2 signaling. (a) Transmitted light images of primary adult rat ventricular myocytes (ARVMs) treated with BSA (control), and mouse recombinant FGF21 (25 ng/ml) with or without 10 mM increased glucose (final 5.6 or 15.6 mM) for 48 hours (scale bar = 25 µm), and (b-d) myocyte length, width, and width to length ratio. FGFR4-specific blocking antibody (anti-FGFR4; 10 mg/ml) was included as indicated. (e) Images of ARVMs treated with BSA (control), and FGF21 (25 ng/ml) with or without phenylephrine (PE; 20 µM) or isoproterenol (Iso; 10 µM) for 48 hours (scale bar = 25 µm), and (f-h) myocyte length, width, and width to length ratio. (i-k) Length, width and width to length ratio for ARVMs treated with BSA (control), FGF21 (25 ng/ml), increased glucose and/or the MEK inhibitor PD98059 (20 µM) for 48 hours. Bars and colored symbols indicate average mean and means of independent experiments using different myocyte preparations, respectively. (l) Binding of 1 µg of soluble β-klotho or PBS, 500 ng of Fc-tagged FGFR 1c, 2c, 3c, or 4 to 96-well plates coated with 200 ng of FGF21. (m) qRT-PCR for FGFR4 mRNA using total RNA isolated from ARVMs treated with BSA (control), FGF21 (25 ng/ml), and/or increased glucose (15.6 mM total). Comparison between groups was performed in form of a one-way (b-d) or two-way (f-k) matched ANOVA followed by post-hoc Tukey test. All values are expressed as mean ± SEM. b-d) N=5, **^**p≤ 0.05 vs. Control CTR, #p≤ 0.05 vs. FGF21 CTR, *p≤ 0.05 vs. Glucose Control. f-h) N=4, *p≤ 0.05 vs. respective CTR. i-k) N=3, **^**p≤ 0.05 vs. BSA CTR, #p≤ 0.05 vs. Glucose CTR, *p≤ 0.05 vs. Glucose +FGF21 CTR; l) N=4, *p≤ 0.05 vs. CTR.

We next considered that FGF21 might drive concentric remodeling in the presence of other stress stimuli. The α-adrenergic agonist phenylephrine induces a selective growth in width of cultured ARVM, while the β-adrenergic agonist isoproterenol induces growth in both width and length, resulting in a more symmetric ARVM hypertrophy (Li et al., 2020a). Co-treatment with FGF21 did not significantly further increase the prominent growth in width induced by phenylephrine or isoproterenol (Figure 2E-H). Remarkably, however, co-treatment with FGF21 inhibited the growth in length induced by isoproterenol, resulting in an increased width:length ratio comparable to that induced by phenylephrine and FGF21. These findings suggest that in the presence of hypertrophic stimuli, including elevated glucose levels, FGF21 will promote a “concentric” phenotype characterized by preferential growth in width of the cardiac myocytes.

Concentric cardiac myocyte hypertrophy can be induced by activation of the ERK1/2 signaling pathway (Kehat et al., 2011). Addition of the MEK inhibitor PD98059 significantly inhibited the growth in width and increased width:length ratio of myocytes cultured in the presence of both FGF21 and high glucose (Figure 2I-K). ERK1/2 signaling is induced by FGF21 in cells expressing β-Klotho and FGFRs (Fisher and Maratos-Flier, 2016), and β-klotho (*Klb*) mRNA was readily detectable in ARVMs (Supplemental Table 3). In a binding assay using recombinant proteins, we found that in the presence of β-klotho, FGF21 bound the Fc-coupled ectodomain of FGFR4 preferentially to FGFR1c, while FGFR2c and FGFR3c did not detectably bind FGF21 at all (Figure 2L). Interestingly, *FGFR4* expression in ARVM was induced synergistically by high glucose and FGF21 (Figure 2M), providing a potential mechanism for the permissive effects of high glucose on FGF21 signaling. Importantly, using a blocking antibody for FGFR4 (Grabner et al., 2015), the synergy between glucose and FGF21 in promoting adult myocyte growth in width was found to require FGFR4 activation (Figure 2B-D).

### Diabetic BTBR ob/ob mice develop FGFR-dependent cardiac hypertrophy

As high serum FGF21 levels are present in diabetic cardiomyopathy (Degirolamo et al., 2016) and FGF21 and high glucose directly promoted cardiac myocyte growth in width, we considered that FGF21/FGFR signaling might promote cardiac hypertrophy in this disease state. The genetic ob/ob mouse model lacking leptin is grossly obese and develops elevated blood glucose (Leibel, 2008; Zhang et al., 1994) and FGF21 levels (Hale et al., 2012; Zhang et al., 2008), while exhibiting cardiac hypertrophy at six months of age (Barouch et al., 2003). ob/ob mice in the BTBR genetic background, which develop severe diabetes (Hudkins et al., 2010), were treated at 3 months of age with the pan-FGFR inhibitor PD173074 by daily intraperitoneal (1 mg/kg i.p.) injection for six weeks. At endpoint, the elevated body weight and blood glucose levels in ob/ob mice (Hudkins et al., 2010; Pichaiwong et al., 2013) were modestly reduced by PD173074 treatment, reaching significance only for body weight (Figure 3A,B). Similarly, serum FGF21 levels were increased 15-fold in BTBR ob/ob mice compared to wild-type littermates and not significantly altered by PD173074 treatment (Figure 3C). BTBR ob/ob mice developed cardiac hypertrophy without fibrosis (Supplemental Figure 2E,F), as indicated by increased LV wall thickness (Figure 3D,E) and cardiac myocyte cross-section area (Figure 3F,G). Notably, the ob/ob-associated hypertrophy was inhibited significantly by PD173074 treatment. These results show that inhibition of FGFR signaling can diminish the development of cardiac hypertrophy despite minimally affecting hyperglycemia and obesity.

**Figure 3.**
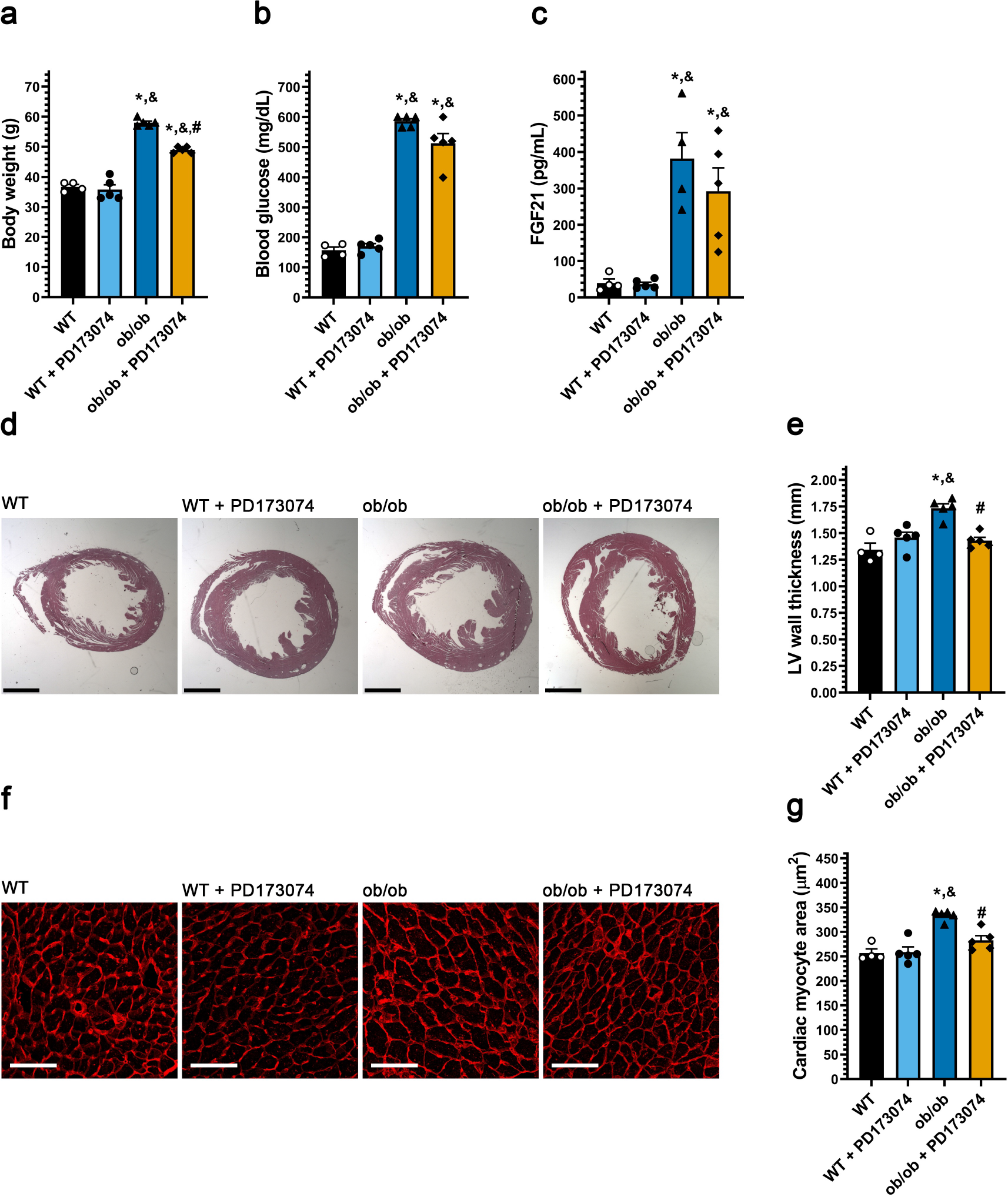
Systemic pan-FGFR blockade protects diabetic mice from cardiac hypertrophy. Three-month old ob/ob and wildtype mice were injected daily with 1 mg/kg PD173074 or vehicle solution for six weeks followed by endpoint analysis. (a) Body weight, (b) blood glucose levels, and (c) serum FGF21 levels. (d) Representative images of H&E-stained transverse heart sections (scale bar = 2 mm) and (e) quantification of the thickness of the LV wall in these images. (f) Wheat germ agglutinin-stained LV tissue (scale bar = 25 µm), and (g) quantification of cardiac myocyte area. Comparison between groups was performed in form of a one-way ANOVA followed by a post-hoc Tukey test (a-c, e, g). All values are expressed as mean ± SEM. a-c, e, g) N=4-5, *p≤ 0.05 vs. WT, &p≤ 0.05 vs. WT+ PD173074 of same age, #p≤ 0.05 vs. ob/ob of same age.

### FGFR4 blockade attenuates cardiac hypertrophy in diabetic db/db mice

*In vitro* data suggested that FGFR4 is the primary FGFR isoform transducing hypertrophic FGF21 signals (Kurosu et al., 2007). This hypothesis was tested *in vivo* using the FGFR4 blocking antibody, in this case using db/db mice (Coleman, 1978), which lack the leptin receptor and develop a more severe cardiomyopathy that includes interstitial myocardial fibrosis. db/db mice, whether saline-injected controls or injected i.p. biweekly with FGFR4 blocking antibody (25 mg/kg) for 24 weeks starting at 4 weeks of age, exhibited elevated body weight, blood glucose and serum FGF21 levels (Figure 4A-C). Notably, the progressive increase in LV wall thickness and LV mass in db/db mice was prevented by anti-FGFR4 treatment (Figure 4D,E, S Supplemental Figure 1C). In addition, the increase in concentricity index at endpoint in db/db mice (Supplemental Table 3), indicative of concentric cardiac hypertrophy, was prevented by treatment with the FGFR4 antibody. Gravimetric analysis confirmed that the blocking antibody inhibited db/db-induced cardiac hypertrophy (Figure 4F). Histologically, db/db mice had increased cardiac myocyte cross-sectional area (Figure 4G,H), as well as interstitial myocardial fibrosis (Supplemental Figure 2G-J), that were both attenuated by the FGFR4 antibody. These results demonstrate that in diabetic db/db mice, FGFR4 activation promotes pathological cardiac remodeling, including concentric hypertrophy and interstitial fibrosis, providing further evidence for a critical role for FGF21-FGFR4 signaling in the cardiomyopathy associated with T2D.

**Figure 4.**
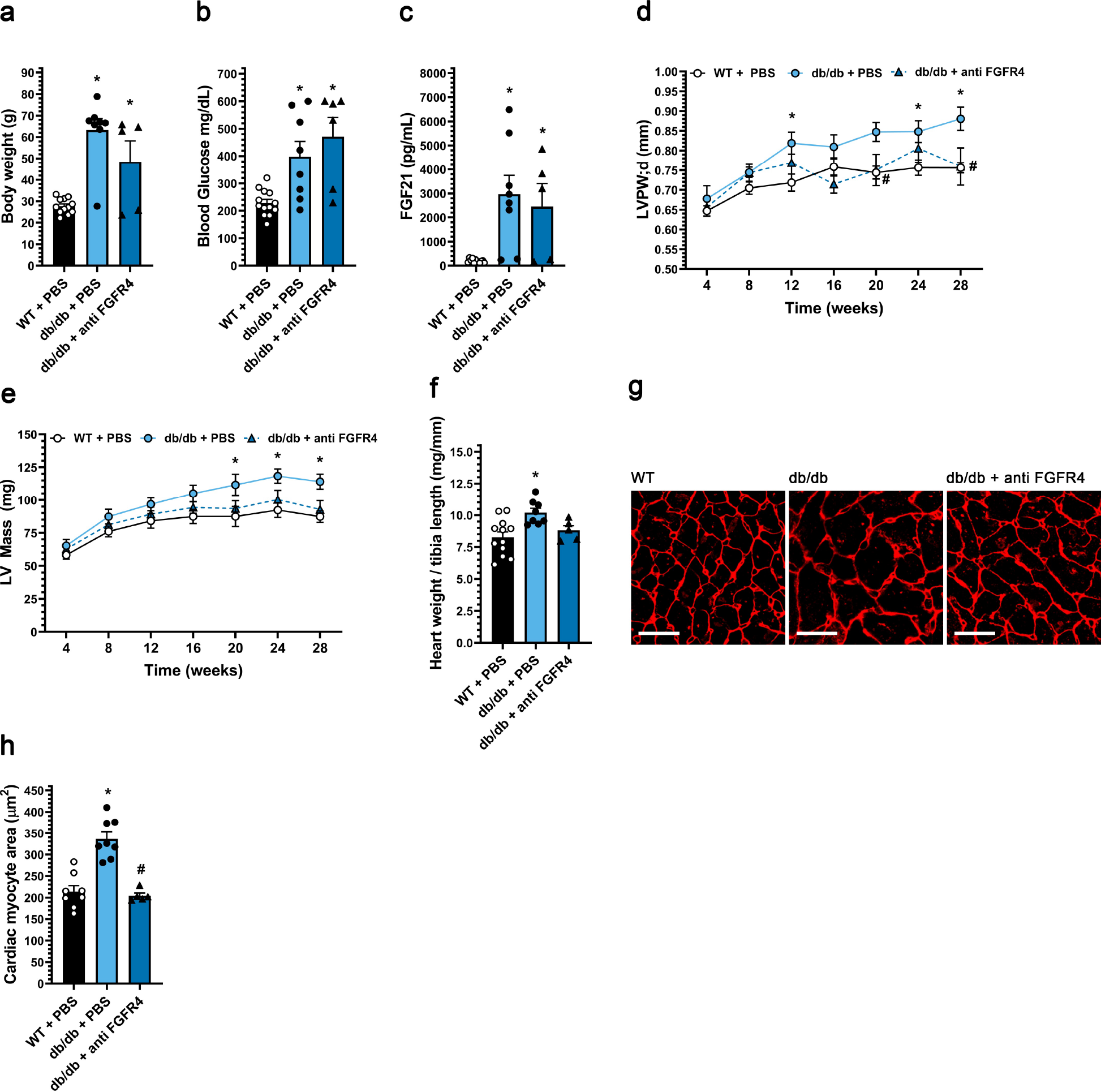
FGFR4 blockade protects diabetic mice from pathologic cardiac remodeling. db/db mice and wild-type (WT) littermates were treated for 24 weeks with either FGFR4 blocking antibody (25 mg/kg) or vehicle (PBS) on a bi-weekly basis, starting at 4 weeks of age. (a) Body weight, (b) blood glucose levels, and (c) serum FGF21 levels. (d) left ventricular (LV) posterior wall thickness in diastole, and (e) LV mass from 4 weeks until 28 weeks of age, as determined by serial echocardiography. (f) Gravimetric heart weight/tibia length ratio, (g) wheat germ agglutinin-stained LV tissue section (scale bar = 25 µm), and (h) cardiac myocyte cross-section area. Comparison between groups was performed in form of a one-way ANOVA (a-c, f, h) or a 2-way ANOVA (d, e) followed by a post-hoc Tukey test. All values are expressed as mean ± SEM. a, c, f) N=5-12; b) N=6-13; h) N=5-9, *p≤ 0.05 vs. WT, #p≤ 0.05 vs. db/db. D, e) N=6-13, *p≤ 0.05 vs. WT of same age, #p≤ 0.05 vs. db/db of same age. For the complete set of echocardiography parameters see Supplemental Table 3.

## Discussion

The development of concentric cardiac hypertrophy is a key feature of the cardiomyopathy associated with diabetes and HFpEF (Mishra and Kass, 2021). Here we provide evidence that elevation in systemic FGF21 levels and the associated cardiac myocyte signaling constitutes a previously unrecognized mechanism for the induction of pathological concentric remodeling, thereby defining a new candidate target for therapeutic intervention in T2D. Blockade of FGFR signaling using both a pan-FGFR small chemical inhibitor and a FGFR4-specific blocking antibody inhibited the concentric hypertrophy associated with diabetic cardiomyopathy in mice. In addition, inhibition of FGFR4 signaling using the blocking antibody and a MEK inhibitor prevented the selective growth in width of adult myocytes *in vitro* induced synergistically by high glucose concentrations and FGF21. Thus, both gain and loss of FGF21-FGFR4 signaling in mouse models, as well as *in vitro* studies using adult cardiac myocytes support in a model in which elevated FGF21 signaling promotes pathological concentric cardiac hypertrophy. Taken in the context of previously published human studies showing elevated FGF21 levels in patients with metabolic syndrome and HFpEF (Chou et al., 2016; Fisher and Maratos-Flier, 2016), we proposed that FGF21 signaling is a conserved critical mechanism contributing to pathological concentric hypertrophy and heart failure.

We found that FGF21 induces myocyte hypertrophy via FGFR4. While others have shown that FGF21 could bind FGFR4-β-Klotho heterodimers (Ogawa et al., 2007), as we show here, it has been suggested that FGF21 cannot activate downstream signaling via FGFR4 (Kurosu et al., 2007; Wu et al., 2010). Instead, we found using that in adult cardiac myocytes, induction of myocyte hypertrophy by FGF21 could be inhibited by a FGFR4-specific blocking antibody. FGF21 is not the only member of the FGF family that can induce cardiac hypertrophy via ERK1/2 signaling. For example, presumably through activation of FGFR1, the paracrine factor FGF2 promotes cardiac hypertrophy under conditions of excess catecholamine stimulation and renin-angiotensin system activation (Itoh and Ohta, 2013). In addition, we have shown that FGF23-FGFR4 signaling can induce cardiac hypertrophy in renal disease (Grabner et al., 2017; Leifheit-Nestler et al., 2016). However, of the endocrine FGFs, the association between elevated FGF21 systemic levels and diabetes and obesity is most well established (Degirolamo et al., 2016). Whether FGF23 is increased in human obesity is controversial (Kutluturk et al., 2019), and we found that FGF23 was not elevated in either the ob/ob or db/db mouse (data not shown). In addition, FGF19 levels are reduced in human obesity (Degirolamo et al., 2016). Thus, while FGF23 may be an important mediator of cardiac hypertrophy in chronic kidney disease, FGF21 may have an especially important role in the induction of concentric hypertrophy in diabetic cardiomyopathy.

The identification of FGF21 as a mediator of pathological concentric hypertrophy stands in contrast to a literature purporting the beneficial effects of FGF21 on the cardiovascular system. While not excluding other mechanisms, including indirect systemic effects, many of these differences may be explained by ERK1/2-mediated differential regulation of concentric and eccentric cardiac hypertrophy and ERK1/2-mediated cardioprotection (Kehat et al., 2011; Lips et al., 2004). As observed *in vitro* for isoproterenol-treated myocytes, FGF21 may promote concentric, but oppose eccentric hypertrophy *in vivo*, regulating the balance between myocyte growth in width and length like ERK1/2 signaling (Kehat et al., 2011). Both ERK1/2 activation by expression of a constitutive MEK1 transgene and ERK1/2 (*Mapk1/Mapk3*) double knock-out resulted in cardiac hypertrophy; however, the former results in a non-fibrotic concentric hypertrophy with thickened myocytes (Bueno et al., 2000), while the latter resulted in ventricular dilation and elongated myocytes (Kehat et al., 2011). This model can explain the exaggerated eccentric hypertrophy in *Fgf21* targeted mice subjected to chronic isoproterenol infusion that promotes eccentric hypertrophy (Planavila et al., 2013). Likewise, the absence of fibrosis in wildtype mice given pharmacologic doses of FGF21 phenocopies MEK1 transgenic mice (Bueno et al., 2000).

Presumably via different downstream effectors that regulate myocyte apoptosis, MEK1 transgenic mice and ERK2 hemizygous knock-out mice exhibit decreased and increased infarct size, respectively, after ischemia-reperfusion (Lips et al., 2004). Likewise, gain and loss of ERK1/2 signaling may explain the beneficial and deleterious effects on myocyte survival and infarct size of exogenous FGF21 and *Fgf21* knock-out, respectively, in mice subjected to ischemia-reperfusion injury or permanent coronary artery ligation (Joki et al., 2015; Liu et al., 2013), although other ERK1/2-independent pathways may also be involved (Hu et al., 2018; Li et al., 2020b; Sun et al., 2019). This pro-survival signaling could complicate the targeting of FGF21-FGFR4 signaling in diabetic cardiomyopathy. However, while *Fgf21* knock-out increased myocyte apoptosis in streptozotocin-induced diabetic mice, *Fgf21* knock-out did not further worsen cardiac dysfunction (Zhang et al., 2015).

The identification of FGF21-FGFR4 signaling as a key regulator of concentric cardiac hypertrophy might provide a novel therapeutic opportunity to prevent or treat heart failure in diabetes. Current cardiac therapeutics have modest or no efficacy for T2D and HFpEF (Falcao-Pires and Leite-Moreira, 2012; Mishra and Kass, 2021). While concentric cardiac hypertrophy can in theory reduce ventricular wall stress (Law of LaPlace), left ventricular hypertrophy is a major risk factor for future heart failure, such that extensive preclinical evidence suggests that preventing concentric hypertrophy even in the presence of persistently increased afterload (hypertension) would be ultimately beneficial (Schiattarella and Hill, 2015). In the context of diabetic cardiomyopathy, concentric hypertrophy predisposes the heart to significant interstitial fibrosis and diastolic dysfunction (Falcao-Pires and Leite-Moreira, 2012). As shown here, inhibited FGFR4 signaling attenuated both cardiac hypertrophy and interstitial myocardial fibrosis in db/db mice. Similarly, inhibition of FGFR4 is a potential approach that might be useful for the cardiomyopathy associated with chronic kidney disease (Grabner et al., 2017). Besides the potential use of biologic agents such as blocking antibodies, there are several recently developed small molecule FGFR4-specific inhibitors being considered for cancer indications in patients (Levine et al., 2020). Alternatively, FGF21 neutralizing antibodies might be developed to inhibit FGF21 function in diabetic cardiomyopathy. Whether non-cardiac effects of FGF21-FGFR4 inhibition would be tolerated in the context of a chronic cardiovascular therapy will determine the potential of this approach. The lack of a severe phenotype in mice with global *Fgfr4* deletion is encouraging (Srisuma et al., 2010; Weinstein et al., 1998). Regardless, readily reversible therapies would be desirable given the beneficial effects of FGF21 in ischemic heart disease, especially since those with diabetes are at risk for a coronary event.

Whether or not FGF21-FGFR4 signaling will prove to be a successful target for intervention against pathological concentric hypertrophy, the observation that FGF21 promotes concentric cardiac hypertrophy should be considered by the community actively developing FGF21 analogs and mimetics as agents to promote weight loss, hyperglycemia and dyslipidemia (Degirolamo et al., 2016). Besides the potential for bone loss, as well as the possibility that there may be “resistance” to the metabolic effects of elevated FGF21 in T2D (Geng et al., 2020), further elevation in FGF21 systemic activity may exacerbate the development of diabetic cardiomyopathy and/or heart failure with preserved ejection fraction.

## Materials and Methods

### Antibodies and Recombinant Proteins

We used human FGF21 (2539-FG-025/CF, R&D Systems, United States), mouse FGF21 (8409-FG-025/CF, R&D Systems), human soluble β-klotho (5889-KB-050/CF, R&D Systems), human FGFR1c (658FR050); human FGFR2c (712FR050); human FGFR3c (766FR050); and human FGFR4 (685FR050). Anti-FGFR4 (human monoclonal, U3-11) was isolated by U3 Pharma/Daiichi-Sankyo (Germany) as described before (Bartz et al., 2019). This antibody is specific for FGFR4 and does not inhibit the activation of FGFR1-3 (Grabner et al., 2015). We used a horseradish peroxidase-conjugated anti-human antibody (109035098, Jackson Immunolabs) for the plate-based binding assay.

### Isolation and Morphometry of Adult Rat Ventricular Myocytes

Eight week-old male Sprague-Dawley rats were anesthetized with Ketamine (40-100 mg/kg i.p.) and Xylazine (5-13 mg/kg i.p.). Hearts were extracted and perfused using a Langendorff system with calcium depletion perfusion buffer containing 120 mM NaCl, 5.4 mM KCl, 1.2 mM NaH2PO4·7H2O, 20 mM NaHCO3, 1.6 mM MgCl2.6H2O, 5 mM taurine, and 5.6 mM glucose, 10 mM 2,3-butanedione monoxime (BDM), followed by type II collagenase digestion at 37°C. The ventricular tissue was cut using sterilized scissors, and the cells were disassociated by pipetting and then filtered using 200 µm mesh. Ventricular myocytes (ARVMs) were collected by centrifugation at 50 *g* for 1 minute. ARVMs were then washed and purified using perfusion buffer containing 0.1% fatty acid-free BSA, 1 mM ATP (pH 7.2), 1 mM glutamine with CaCl2 added gradually until reaching a Ca^2+^ concentration of 1.8 mM. The purified ARVMs were then resuspended in a defined Minimal Medium [Medium 199 (11150059, Thermo Fisher, United States) supplemented with 5 mM creatine, 2 mM L-carnitine, 5 mM taurine, 25 mM HEPES, 0.2% fatty acid-free BSA, 10 mM BDM and insulin-transferrin-selenium (ITS)] and seeded onto square cover slips (21 mm x 21 mm) coated with mouse laminin (10 μg/mL diluted in PBS) in 6-well plates (10,0000 cells per well). One hour after isolation, the culture medium was removed, ARVMs were washed with 2 mL medium once, then cultured for 48 hours in Minimal Medium containing recombinant FGF21 (25 ng/mL, dissolved in 0.1% BSA) or 0.1% BSA control with and without phenylephrine (20 μM), isoproterenol (10 μM) or additional glucose (10 mM, final 15.6 mM) as indicated. To block FGFR4, isolated ARVMs were pre-treated with Minimal Medium containing FGFR4 blocking antibody (10 mg/ml human monoclonal U3-11; U3Pharma) for one hour before addition of FGF21 (25 ng/mL) and glucose (15.6 mM final). For morphometric analysis, ARVMs were washed twice with PBS and fixed with 3.7% formaldehyde solution (diluted in PBS) for one hour. Cover slips were mounted onto slides, and nine images per slide were obtained using a Leica DM4000 Microscope. The width and length of 100 ARVMs/image were measured by Leica LAX software. ARVM length and width were defined by the maximum dimensions of the cell either parallel and perpendicular to sarcomeric striations, respectively.

### Plate-Based Binding Assay

To measure FGF21 binding to FGFRs, 96-well plates (Thermo Fisher; #44-2404-21) were coated with 200 ng of human FGF21 protein in 100 µL of coating buffer (E107, Bethyl) per well at 4°C overnight. Plates were washed 5x with 350 µL of assay buffer (50 mM Tris pH 7.4, 200 mM NaCl, 0.01% Tween 20) on a 50TS microplate washer (BioTek, United States). Plates were blocked for 1 hour in 200 µL assay buffer with 0.5% BSA (Rockland; BSA-50). Plates were washed 5x with assay buffer and incubated with 100 µL assay buffer with 0.5% BSA in combination with PBS or 1 µg of human soluble β-klotho at room temperature. After 1 hour, plates were washed, and 500 ng of FGFR-Fc in 100 µL volume of assay buffer with 0.5% BSA were added per well. After 1 hour at room temperature, plates were washed 5x in assay buffer and incubated with 100 µL anti-human Fc-HRP at 1:10,000. Plates were washed as above and 100µL TMB substrate (E102, Bethyl) was added for 15-20 minutes until positive wells developed a dark blue color. Reactions were stopped with an ELISA stop solution (E115, Bethyl) and analyzed on a Synergy H1 plate reader (BioTek) at 450 nm wavelength. All samples were run in triplicates

### RNA Isolation and Real-Time PCR

RNA from total mouse hearts was extracted using Trizol (15596026, Invitrogen, United States). 2 µg total RNA was reverse transcribed using Qscript (95048, Quanta Biosciences, United States). Quantitative PCR reactions were carried out in the StepOne plus Real-Time PCR System (Applied Biosystems, United States) using FAST SYBR Green Master Mix (4385610, Applied Biosystems). Raw data was quantified via StepOneTM software v2.3 from life Technologies. Relative gene expression was normalized to expression levels of GAPDH and evaluated using the 2-ΔΔCt method.

ARVM RNA was extracted using RNeasy mini kit (74104, Qiagen, United States) according to manufacturer’s instructions. 500 ng total RNA was reverse transcribed using a Reverse Transcription Kit (4368814, Applied Biosystems). Quantitative PCR reactions were performed using the QuantStudio 5 System (Thermo Scientific, United States) using Powerup SYBR Green Master Mix (A25742, Thermo Scientific). Raw data was analyzed using QuantStudio 5 qPCR Data Analysis Software (Thermo Scientific). Relative gene expression was normalized to expression levels of Hprt1 and evaluated using the 2-ΔΔCt method. Primer sequences are presented in Table 3.

### FGF21 Transgenic Mice

FGF21 transgenic mice expressing a mouse FGF21 cDNA under the control of the apolipoprotein (ApoE) promoter (C57BL/6-Tg(Apoe-Fgf21)1Sakl/J; The Jackson Laboratory Stock No: 021964) were as previously described (Inagaki et al., 2007). Eleven hemizygous and 10 wild type mice, male and female, were analyzed by echocardiography at 4 and 6 months of age before euthanasia for further analysis.

### FGF21 Serial Injections

FGF21 injections were conducted following a similar protocol as previously established for FGF23 by us (Faul et al., 2011) and others (Andrukhova et al., 2014; Han et al., 2019). Briefly, 12-week-old, male and female BALB/cJ mice (Stock No: 000651; The Jackson Laboratory, United States) underwent tail vein injections. The day before the experiment, mice underwent echocardiographic analysis, as described below. Mice were anesthetized using 2.5% isoflurane and placed on a heat pad. Mice were divided into 2 groups of 10, receiving either vehicle solution (isotonic saline) or mouse FGF21 protein. Per injection, we used 40 µg/kg of FGF21 dissolved in 200 µL of isotonic saline, with 8 hours between injections for a total of 5 consecutive days. All injections were performed via the lateral tail vein. On the morning of the sixth day, 16 hours after the final tail vein injection, animals underwent echocardiographic analysis and were sacrificed.

### ob/ob mice

Ob/ob mice lacking leptin (Zhang et al., 1994) were studied in the genetic BTBR background (BTBR.Cg-*Lep^ob^*/WiscJ; Stock No: 004824; The Jackson Laboratory) (Hudkins et al., 2010). At 3 months of age, 5 ob/ob and 5 wild type mice, male and female, were i.p. injected daily with PD173074 (P2499, Sigma Aldrich) at 1 mg/kg and 5 ob/ob and 4 wild type mice with saline for 6 weeks. Animals were sacrificed and samples prepared as described here.

### db/db mice

Db/db mice which lack the leptin receptor (Coleman, 1978) were purchased from The Jackson Laboratory (B6.BKS(D)-*Lepr^db^*/J; Stock No: 000697). Five 4-week-old db/db mice were i.p. injected bi-weekly with 25 mg/kg of anti FGFR4 for 24 weeks. Eight db/db mice and 13 wild type mice were injected with vehicle as control. All groups contained male and female mice. Animals were monitored every 4 weeks for body weight and blood glucose levels and by echocardiographic measurements. After 24 weeks, animals were sacrificed, and samples were prepared as here.

### Histology and Morphometry of Mouse Hearts

Heart tissue was fixed overnight in 4% phosphate-buffered formaldehyde solution. Tissue was sent to IDEXX Bioanalytics (Columbia, MO) for embedding and sectioning. Transverse sections stained with hematoxylin and eosin (H&E) were imaged on a Leica DMi8 microscope.

We measured the cross-section area of individual cardiac myocytes in paraffin-embedded transverse sections. Paraffin sections underwent deparaffinization 2x for 5 minutes in Shandon Xylene Substitute and then rehydrated through a graded ethanol series (99%, 97%, 70%), 2x for 5 minutes each. Antigen retrieval was performed in a microwave for 15 minutes in 1x unmasking solution (H3300, Vector Labs). Slides were washed 3x for 5 minutes each in PBS, then incubated for 1 hour in blocking solution ((1% BSA50 (Rockland), 0.1% cold water fish skin gelatin (900033, Aurion), and 0.1% Tween 20)). Slides were washed 3x in PBS and then incubated in 10 µg/mL of 594-conjugate WGA (W11262, Thermo Fisher) for 1 hour. Slides were washed 3x with PBS and then mounted in Prolong Diamond (P36961, Thermo Fisher). Immunofluorescence images were taken on a Leica DMi8 microscope with a 63x oil objective. ImageJ software (NIH) was used to quantify the cross-section area of 25 myocytes per field in 4 fields along the mid-chamber free wall based on WGA-positive staining.

### Echocardiography of Mouse Hearts

For mice receiving serial injections of recombinant FGF21 protein, transthoracic echocardiographic analysis was performed on day 6 of the experiment, at 13 weeks of age using a Vevo 770® High-Resolution Micro-Imaging System (FUJIFILM VisualSonics), equipped with an 707B-253 transducer. Animals were minimally anesthetized with 1-1.5% isoflurane, and normal body temperature was maintained using an anal probe for temperature monitoring. For analysis, both B- and M-mode images were obtained in the short and long axis view. Correct positioning of the transducer was ensured using B-mode imaging in the long axis view, before switching to the short axis view. Parameters were calculated from at least three cardiac cycles by M-mode echocardiographs. All parameters were measured or calculated using Vevo® 770 Workstation Software (FUJIFILM VisualSonics).

Serial transthoracic echocardiography was performed in FGF21 Tg mice at the age of 16 and 24 weeks, and in db/db mice every 4 weeks from 4 weeks of age, using a Vevo 2100 High-Resolution Imaging System (FUJIFILM VisualSonics). Animals were minimally anesthetized with 1-1.5% isoflurane, and normal heart rate and body temperature were monitored and maintained throughout the procedure. Parameters were calculated from at least three cardiac cycles by M-mode echocardiographs. All parameters were measured or calculated using the VevoLAB software (FUJIFILM VisualSonics). Measurement definitions of short axis and calculations (M-Mode) are presented in Table 1 and 2.

**Table 1.** Measurement definitions short axis (M-Mode)

| **Parameter** | **Definition** | **Unit** |
| --- | --- | --- |
| LVPW;d | Left ventricular posterior wall<br>(diastole) | mm |
| LVPW;s | Left ventricular posterior wall<br>(systole) | mm |
| LVAW;d | Left ventricular anterior wall<br>(diastole) | mm |
| LVAW;s | Left ventricular anterior wall<br>(systole) | mm |
| LVID;d | Left ventricular internal<br>diameter (diastole) | mm |
| LVID;s | Left ventricular internal<br>diameter (systole) | mm |
| Heart Rate | Heart Rate | BPM |
| Body Weight | Body Weight | g |
| LV Mass/Body weight | LV Mass/Body weight ratio | mg/g |

**Table 2.** Calculations short-axis (M-Mode)

| Parameter | Definition | Unit | Formula |
| --- | --- | --- | --- |
| LV Vol;d | Left ventricle volume diastole (M-Mode) | μl | $\left(\frac{7.0}{2.4 + LVID; d}\right) \times LVID; d^3$ |
| LV Vol;s | Left ventricle volume systole (M-Mode) | μl | $\left(\frac{7.0}{2.4 + LVID; s}\right) \times LVID; s^3$ |
| EF | LV ejection fraction (M-Mode) | % | $100 \times \left(\frac{LV Vol; d - LV Vol; s}{LV Vol; d}\right)$ |
| FS | LV fractional shortening (M-Mode) | % | $100 \times \left(\frac{LVID; d - LVID; s}{LVID; d}\right)$ |
| LV Mass AW (Corrected) | LV Mass AW corrected (M-Mode) | mg | $0.8 \times (1.053 \times ((LVID; d + LVPW; d + LVAW; d)^3 - LVID; d^3))$ |
| Concentricity | | | $\left(\frac{LVAW; d + LVPW; d}{LVID; d}\right)$ |

IVS;d: Inter ventricular septum (diastole) in mm

**Table 3.**
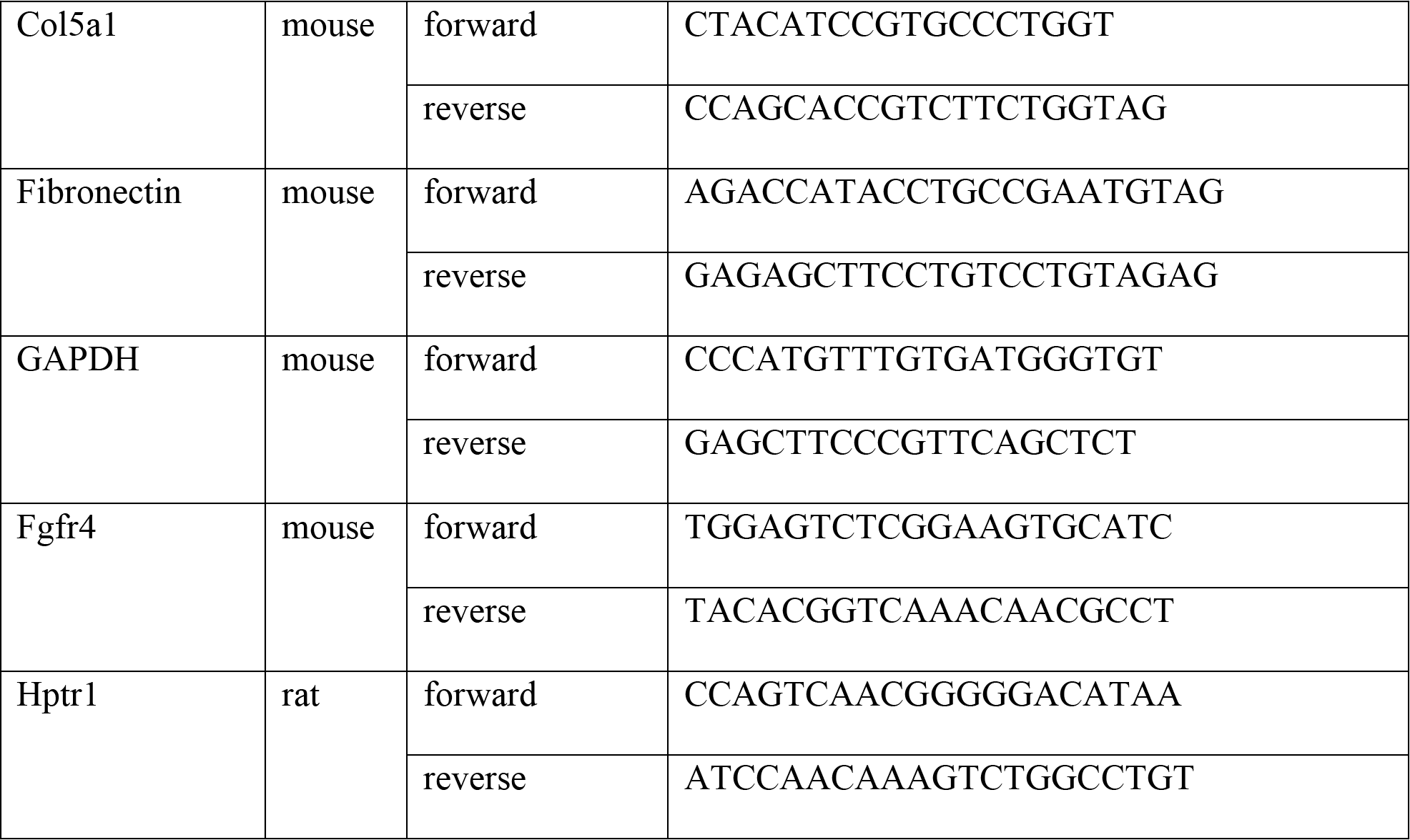
Primer sequences.

### Serum Chemistry

At endpoint, blood was collected from mice via cardiac puncture, transferred into microvette serum gel tubes (20.1344, Sarstedt) and centrifuged at 10,000 g for 5 minutes. Serum supernatants were collected and stored at -80°C. ELISAs were run to detect FGF21 (MF2100, R&D Systems) and FGF23 (60-6300; Quidel).

### Statistics

Values are expressed as mean ± SEM if not otherwise indicated. Comparisons between 3 or more groups were performed by one-way ANOVA followed by post-hoc Tukey test or by 2-way ANOVA with post-hoc Sidak’s multiple comparison test. Comparisons between 2 groups were performed by two-tailed t-tests. A significance level of P≤0.05 was accepted as statistically significant. No statistical method but experience from previous publications was used to predetermine sample size. No formal randomization was used in any experiment. For *in vivo* experiments, animals were unbiasedly assigned into different treatment groups. Group allocation was not performed in a blinded manner. Whenever possible, experimenters were blinded to the groups (for example, in IF and IHC experiments by hiding group designation and genotype of animals until after quantification and analysis).

### Study Approval

All animal protocols and experimental procedures for FGF21 injections in mice, studying FGF21 Tg mice, and pan-FGFR and anti-FGFR4 injections in ob/ob and db/db mice were approved by the Institutional Animal Care and Use Committees (IACUC) at the University of Alabama Birmingham School of Medicine and the University of Miami Miller School of Medicine. The use of rats for myocyte isolation was approved by the Administrative Panel on Laboratory Animal Care at Stanford University. All animals were maintained in temperature-controlled environments with a 12-hour light/dark cycle and allowed *ad libitum* access to food and water. All protocols adhered to the Guide for Care and Use of Laboratory Animals to minimize pain and suffering.

## Acknowledgments

C.Y., D.K., X.L., A.G., M.S.K. and C.F. designed the study and analyzed the data. C.Y. injected protein in mice, and A.G., K.S. and A.S. injected inhibitors in mice. E.C.M. and D.K. conducted echocardiography, and M.S.K. and A.W. assisted with the analysis of echocardiographic data. A.G., K.S. and D.K. conducted histological and serum chemistry analyses of mice. I.C., B.C., K.H. and D.W. assisted with animal studies. X.L. conducted studies in isolated rat cardiac myocytes. A.F. assisted with data interpretation from diabetic mouse models. C.Y., M.S.K. and C.F. wrote the paper, and D.K. designed the figures.

This study was supported by the Deutsche Forschungsgemeinschaft (D.K.), the National Science Foundation (I.C.), the American Society of Nephrology (A.G.), the American Heart Association (E.C.M.), the American Diabetes Association (C.F.), and grants F31DK115074 (C.Y.), F31DK117550 (B.C.), R01HL133011 (A.W.), R01DK117599, R01DK104753, R01CA227493, U54DK083912, UM1DK100846, and U01DK116101 (A.F.), R01HL146111, R01 HL158052 and R01HL126825 (M.K.), R01HL128714 and R01HL145528 (C.F.) from the NIH. A.F. was supported by the Miami Clinical Translational Science Institute (UL1TR000460), and C.F by the UAB-UCSD O’Brien Core Center for Acute Kidney Injury Research, the AMC21 program of the Department of Medicine at UAB, and the Tolwani Innovation Award from the Division of Nephrology at UAB.

## Supplemental Data

**Supplemental Figure 1:**
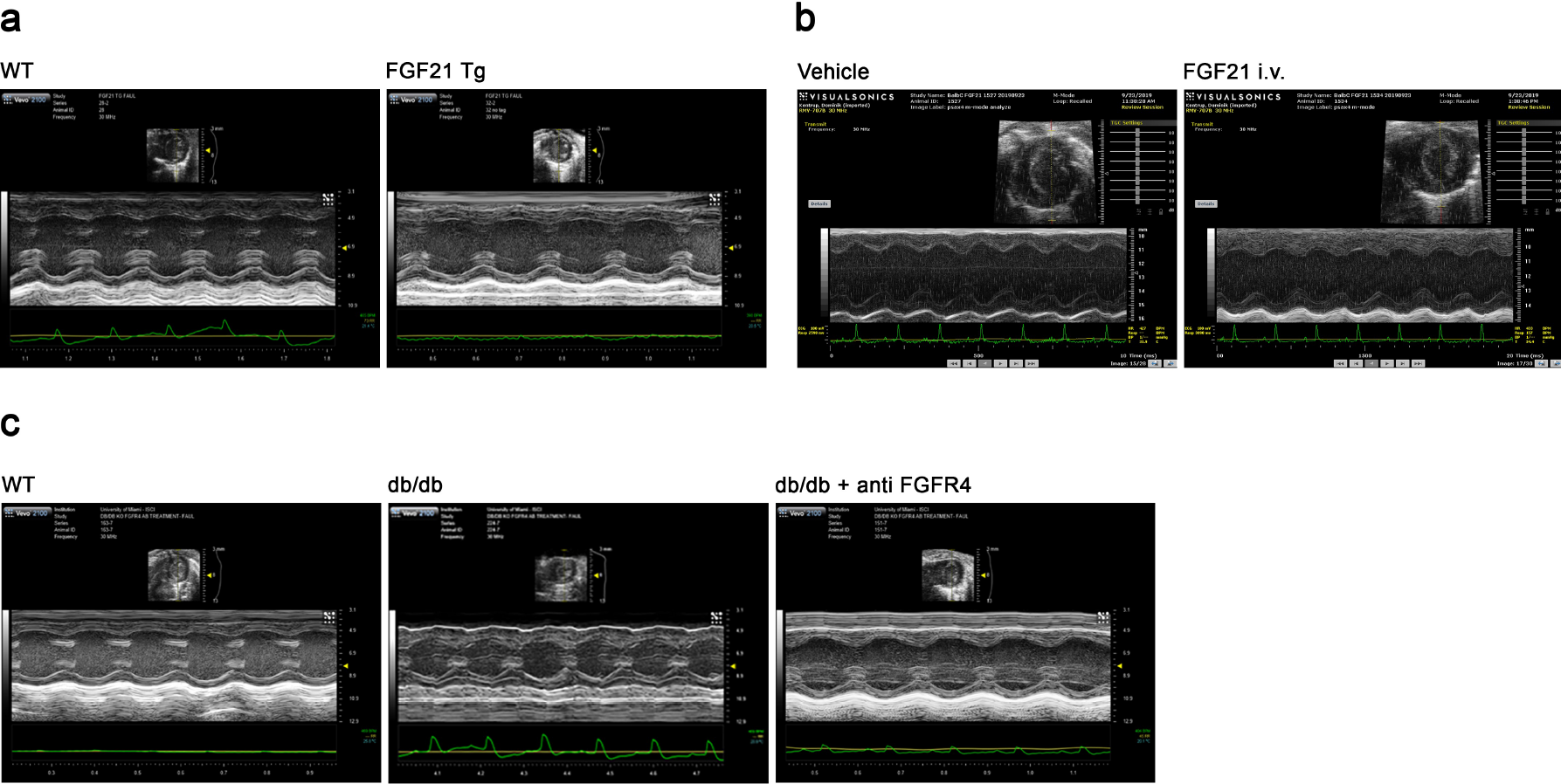
Echocardiography M-mode images (short axis view) Representative M-mode images of (a) FGF21 transgenic (Tg) mice and wild-type littermates at 24 weeks of age, (b) wildtype mice i.v. injected with either FGF21 or vehicle solution for five consecutive days, and (c) db/db mice and wild-type littermates at 28 weeks of age, after being treated for 24 weeks with either anti-FGFR4 (25 mg/kg) or vehicle solution on a bi-weekly basis, starting at 4 weeks of age.

**Supplemental Figure 2:**
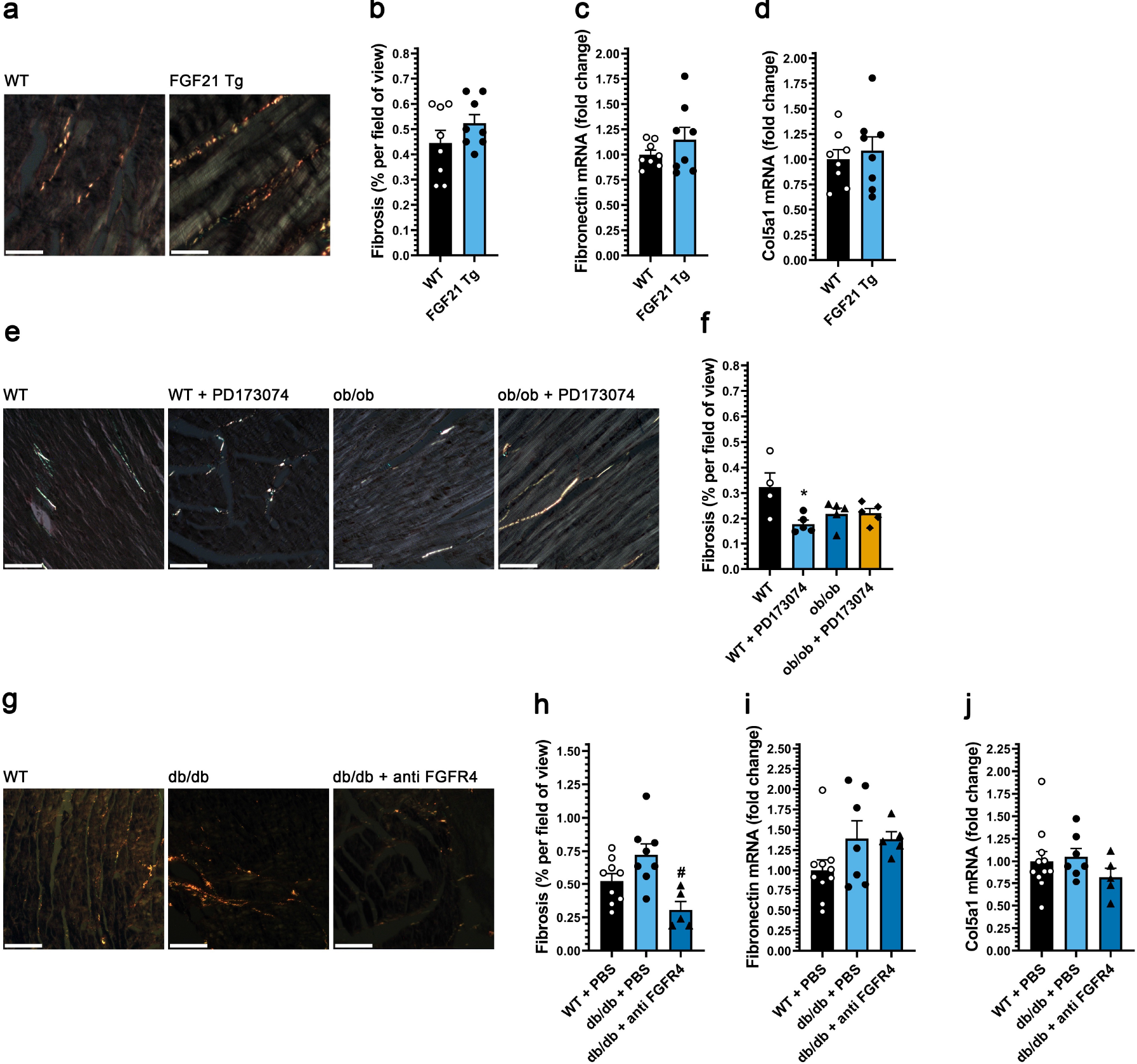
Analysis of cardiac fibrosis. (a) Representative images of cardiac tissue sections of FGF21 transgenic (Tg) mice, stained with picrosirius red, taken under polarized light (scale bar = 50 µm), and (b) quantification of positive areas. Real-time qPCR analysis of fibrosis markers (c) fibronectin and (d) Col5a1 in cardiac tissue. (e) Representative images of cardiac tissue sections stained with picrosirius red, taken under polarized light (scale bar = 50 µm), of ob/ob mice and wild-type littermates at three months of age, after being treated daily with 1 mg/kg PD173074 or vehicle solution for six weeks (scale bar = 2 mm), and (f) the quantification of stained area. (g) Representative images of cardiac tissue sections of db/db mice and wild-type littermates at 28 weeks of age, after being treated for 24 weeks with either anti-FGFR4 (25 mg/kg) or vehicle solution on a bi-weekly base, starting at 4 weeks of age, stained with picrosirius red, taken under polarized light (scale bar = 50 µm), and (h) the quantification of stained area. (i, j) Real-time qPCR analysis of fibrosis markers (j) fibronectin and (k) Col5a1 in cardiac tissue. Comparison between groups was performed in form of a one-way ANOVA (f, h-j) followed by a post-hoc Tukey test, or in form of a two-tailed t-test (b-d). A significance level of p≤0.05 was accepted as statistically significant. All values are expressed as mean ± SEM. b) N=8, *p≤ 0.05 vs. WT; c, d) N=8, *p≤ 0.05 vs. WT; f) N=4-5, *p≤ 0.05 vs. WT; h) N=5-9, *p≤ 0.05 vs. WT, #p≤ 0.05 vs. db/db; i, j) N=5-11, *p≤ 0.05 vs. WT, #p≤ 0.05 vs. db/db.

**Supplemental Table 1:** Echocardiography of FGF21 Tg mice at 16 and 24 weeks of age.

|  |  | **<u>WT</u>**<br>**N=8** | **<u>FGF21 Tg</u>**<br>**N=8** |  |
| --- | --- | --- | --- | --- |
| **<u>16 Weeks</u>** |  |  |  |  |
| LVPW;d | mm | 0.71 ± 0.06 | 0.62 ± 0.08 | * |
| LVPW;s | mm | 0.97 ± 0.09 | 0.97 ± 0.21 |  |
| LVAW;d | mm | 0.64 ± 0.08 | 0.57 ± 0.11 |  |
| LVAW;s | mm | 0.95 ± 0.13 | 0.92 ± 0.15 |  |
| LVID;d | mm | 4.07 ± 0.24 | 3.38 ± 0.26 | *** |
| LVID;s | mm | 2.94 ± 0.30 | 2.09 ± 0.37 | *** |
| LV Volume;d | μL | 75 ± 10 | 48 ± 7 | *** |
| LV Volume;s | μL | 35 ± 7 | 16 ± 6 | *** |
| Ejection fraction | % | 54 ± 7 | 67 ± 9 | *** |
| Fractional shortening | % | 28 ± 4 | 37 ± 7 | ** |
| LV Mass | mg | 85 ± 13 | 49 ± 8 | *** |
| Body weight | g | 26 ± 5 | 14 ± 1 | *** |
| LV Mass/Body weight | mg/g | 3.4 ± 0.5 | 3.5 ± 0.4 |  |
| Heart rate | BPM | 438 ± 30 | 434 ± 32 |  |
| Concentricity |  | 0.33 ± 0.03 | 0.35 ± 0.06 |  |
|  |  | **<u>WT</u>**<br>**N=8** | **<u>FGF21 Tg</u>**<br>**N=8** |  |
| **<u>24 Weeks</u>** |  |  |  |  |
| LVPW;d | mm | 0.74 ± 0.06 | 0.76 ± 0.05 |  |
| LVPW;s | mm | 1.04 ± 0.09 | 1.20 ± 0.11 | * |
| LVAW;d | mm | 0.69 ± 0.09 | 0.75 ± 0.05 |  |
| LVAW;s | mm | 1.03 ± 0.11 | 1.17 ± 0.07 |  |
| LVID;d | mm | 3.98 ± 0.25 | 3.38 ± 0.26 | *** |
| LVID;s | mm | 2.75 ± 0.24 | 2.08 ± 0.20 | *** |
| LV Volume;d | μL | 70 ± 10 | 50 ± 8 | *** |
| LV Volume;s | μL | 29 ± 7 | 15 ± 2 | *** |
| Ejection fraction | % | 58 ± 6 | 70 ± 3 | ** |
| Fractional shortening | % | 31 ± 4 | 39 ± 2 | ** |
| LV Mass | mg | 81 ± 11 | 67 ± 10 | ** |
| Body weight | g | 28 ± 6 | 15 ± 2 | *** |
| LV Mass/Body weight | mg/g | 3.0 ± 0.3 | 4.5 ± 0.6 | *** |
| Heart rate | BPM | 479 ± 47 | 425 ± 36 | * |
| Concentricity |  | 0.36 ± 0.04 | 0.45 ± 0.03 | *** |

Abbreviations: LVPW, Left ventricular posterior wall thickness; LVAW, left ventricular anterior wall thickness; LVID, Left ventricular internal diameter; LV, Left ventricular; d, diastole; s, systole; concentricity = (LVAW;d+LVPW;d)/LVID;d in diastole; BPM, beats per minute. All data are mean ± SD. \*p≤ 0.05, \*\*p<0.01, \*\*\*p<0.001 vs. WT of same age.
All p-values are based upon 2-way ANOVA Sidek post-hoc testing involving both datasets.

**Supplemental Table 2:**
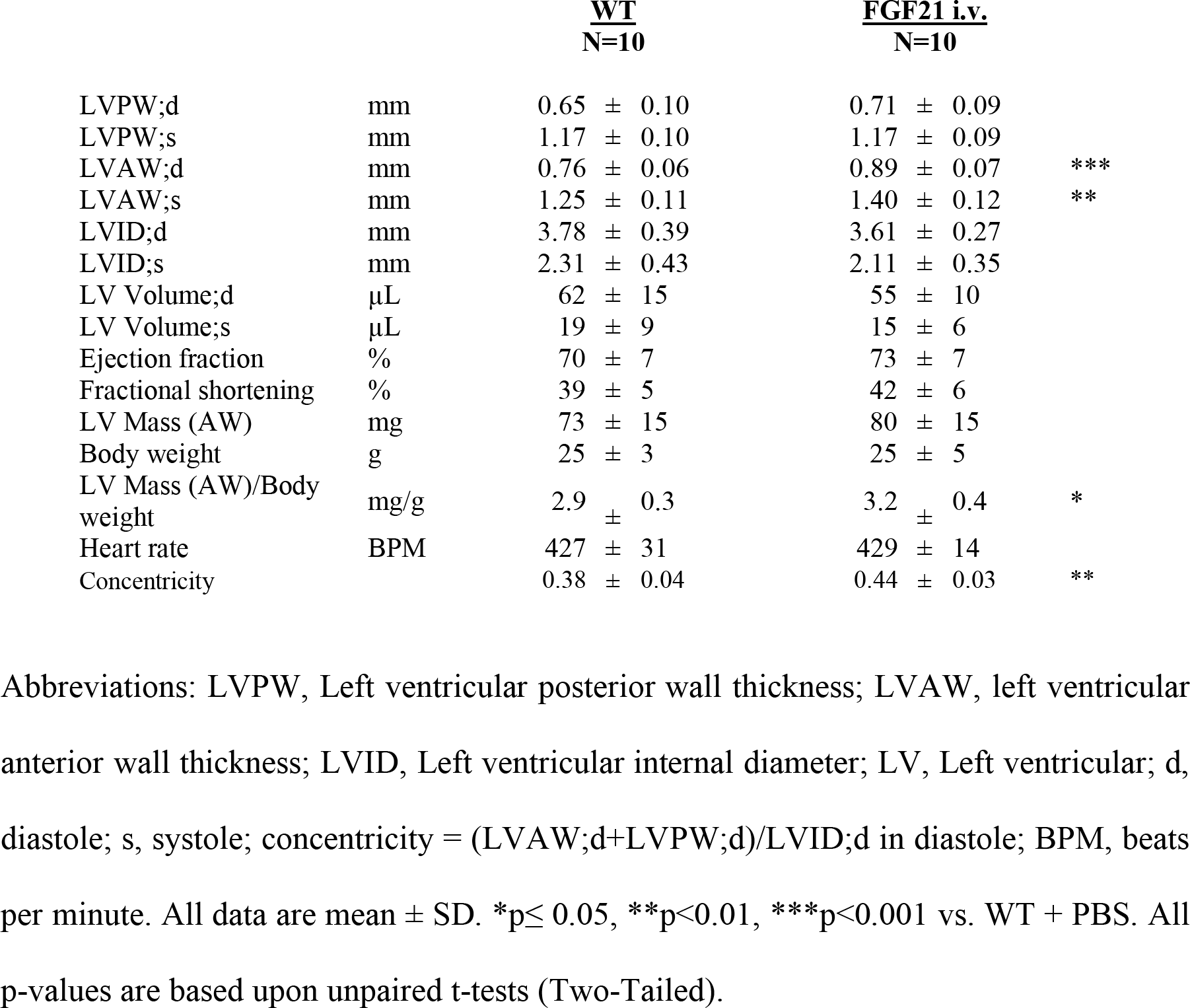
Echocardiography of mice receiving serial FGF21 injections.

**Supplemental Table 3:** Echocardiography of db/db mice receiving anti-FGFR4 for 28 weeks.

|  |  | <u>WT + PBS</u><br>N=13 | <u>db/db + PBS</u><br>N=9 |  | <u>db/db + anti FGFR4</u><br>N=6 |  |
| --- | --- | --- | --- | --- | --- | --- |
| **<u>28 weeks</u>** |  |  |  |  |  |  |
| LVPW;d | mm | 0.76 ± 0.01 | 0.88 ± 0.03 | ** | 0.76 ± 0.05 | # |
| LVPW;s | mm | 1.00 ± 0.02 | 1.26 ± 0.07 | *** | 1.01 ± 0.07 | ## |
| LVAW;d | mm | 0.70 ± 0.01 | 0.91 ± 0.07 | *** | 0.69 ± 0.05 | ### |
| LVAW;s | mm | 1.07 ± 0.03 | 1.36 ± 0.10 | *** | 1.02 ± 0.07 | ### |
| LVID;d | mm | 4.05 ± 0.12 | 4.11 ± 0.17 |  | 4.28 ± 0.13 |  |
| LVID;s | mm | 2.85 ± 0.14 | 2.76 ± 0.22 |  | 2.95 ± 0.22 |  |
| LV Volume;d | μL | 76 ± 5 | 76 ± 7 |  | 84 ± 5 |  |
| LV Volume;s | μL | 34 ± 4 | 32 ± 6 |  | 35 ± 5 |  |
| Ejection fraction | % | 57 ± 2 | 60 ± 4 |  | 58 ± 5 |  |
| Fractional shortening | % | 30 ± 2 | 32 ± 3 |  | 31 ± 3 |  |
| LV Mass | mg | 87 ± 4 | 114 ± 6 | ** | 93 ± 7 |  |
| Body Weight | g | 27 ± 1 | 56 ± 7 | *** | 46 ± 9 | *** |
| Heart rate | BPM | 460 ± 12 | 455 ± 15 |  | 401 ± 23 |  |
| Concentricity |  | 0.36 ± 0.04 | 0.45 ± 0.11 | *** | 0.34 ± 0.07 | ### |

Abbreviations: LVPW, Left ventricular posterior wall thickness; LVAW, left ventricular anterior wall thickness; LVID, Left ventricular internal diameter; LV, Left ventricular; d, diastole; s, systole; concentricity = (LVAW;d+LVPW;d)/LVID;d in diastole; BPM, beats per minute. All data are mean ± SD. \*p≤ 0.05, \*\*p<0.01, \*\*\*p<0.001 vs. WT + PBS of same age, #p≤ 0.05, ##p<0.01, ###p<0.001 vs. db/db + PBS of same age. All p-values are based upon 2-way ANOVA and Tukey post-hoc testing involving full-time course dataset.

